# Organic management promotes natural pest control through enhanced plant resistance to insects

**DOI:** 10.1101/787549

**Authors:** Robert Blundell, Jennifer E. Schmidt, Alexandria Igwe, Andrea L. Cheung, Rachel L. Vannette, Amélie C.M. Gaudin, Clare L. Casteel

## Abstract

Lower insect pest populations found on long-term organic farms have largely been attributed to increased biodiversity and abundance of beneficial predators. However, potential induction of plant defenses has largely been ignored. This study aims to determine whether host plant resistance mediates decreased pest populations in organic systems, and to identify the underpinning mechanisms. We demonstrate that greater numbers of leafhoppers *(Circulifer tenellus*) settle on tomatoes (*Solanum lycopersicum*) grown using conventional management as compared to organic. Soil microbiome sequencing, chemical analysis, and transgenic approaches, coupled with multi-model inference, suggest that changes in leafhopper settling between organically and conventionally-grown tomatoes are dependent on salicylic acid accumulation in the plant, likely mediated by rhizosphere microbial communities. These results suggest that organically-managed soils and microbial communities may play an unappreciated role in reducing plant attractiveness to pests by increasing plant resistance.

---

Organic farming is characterized by management practices that promote biodiversity and beneficial ecological interactions to offset synthetic inputs such as inorganic fertilizers and biocides. Pest and nutrient management in organic agriculture is largely accomplished through diversification methods such as cover crops, crop rotations, and trap crops. Organic agriculture is often thought to be less productive in terms of yield as compared to conventional farming, even though it offers great potential as a more sustainable agricultural system (4, 5, 6) and can enhance the provision of multiple ecosystem services, such as carbon sequestration, nutrient and water retention, and biodiversity conservation (1–3).

Accumulating evidence suggests that organic management practices also reduce pest populations and increase resilience to pest damage (7-9). Enhancing natural pest control in organic systems could help reduce costs, stabilize production, and increase the ability of organic practices to meet global demand. Decreased insect pests on long-term organic farms have largely been attributed to practices that limit pest build-up, increase predator biodiversity, and increase the numbers of beneficial insects (7, 10, 11). It is well-known that herbivore populations are also influenced by plant defense responses, but the impact of organic management on plant defense capacity has been largely ignored.

Organic management strategies can increase microbial activity and biomass in soils (12–14), alter microbial communities, and in some cases enhance plant associations with beneficial microbes in the rhizosphere (2, 15, 16). Microorganisms that associate with plant roots play a critical role in resistance to abiotic and biotic stress (17–19). For example, mycorrhizal fungi have been shown to induce systemic resistance throughout the plant (20, 21) and can reduce susceptibility to pathogens (22) and herbivores (23). Further, plant growth-promoting rhizobacteria (PGPR) commonly found in soil microbial pools as well as commercial inoculants, can induce defenses and other physiological changes in the host plant that influence aboveground herbivores (18, 24–26). Despite the known interactions between organic management, plant-microbe associations, and changes in crop resistance, the potential of these interactions to reduce pest damage in agricultural systems remains largely untapped.

The objectives of this study are to determine whether organic approaches to management influence pest populations through changes in plant resistance and to investigate potential mechanisms. We explore linkages between insect populations, leafhopper *(Circulifer tenellus)* settling and performance, rhizosphere communities, plant nutrients, and phytohormones related to plant defense in tomato (*Solanum lycopersicum*) using on-farm and in-lab studies. The beet leafhopper is routinely found in California tomato fields and it is an important pest due to its ability to transmit Beet curly top virus (BCTV) to tomatoes. In 2013, an outbreak of BCTV resulted in ~$100 million losses for California’s processing tomato industry (27). We demonstrate that tomatoes grown using conventional management are preferentially settled by leafhopper pests and have reduced salicylic acid (SA) levels compared to tomatoes grown using organic management. Differences in insect preference were due partially to changes in SA accumulation and rhizosphere microbial communities. Understanding how soil management influences plant resistance and to what extent it helps create robust and resilient systems will provide growers with new pest management tools to improve multiple sustainability outcomes for agroecosystems.

## Results

### Organic management reduced insect populations and settling on tomatoes

To determine if management influenced plant attractiveness to insects, we collected tomato branches from organic and conventional plots at the long-term experimental farm Russell Ranch (Farm RR) in Davis, CA and at three commercial farms in Yolo county in 2017 (Farm RR, Farm F, Farm M, and Farm S; See Table S1). These branches were used to compare beet leafhopper settling preference for the leaves paired by site (organic versus conventional; See Fig. S1 for design). Fewer leafhoppers settled on tomato leaves from organic sites at three out of the four locations compared to tomato leaves grown on conventional plots (Fig. 1A; Farm RR, Farm M, and Farm S). Next, we surveyed insect populations using sweepnet sampling in the same organic and conventionally managed tomato plots. We observed significantly fewer insects in organic plots compared to conventional plots at Farm RR (Fig. S2). No systematic differences in insect abundance were observed between organic and conventional plots at the other sites (Fig. S1B).

**Fig. 1.**
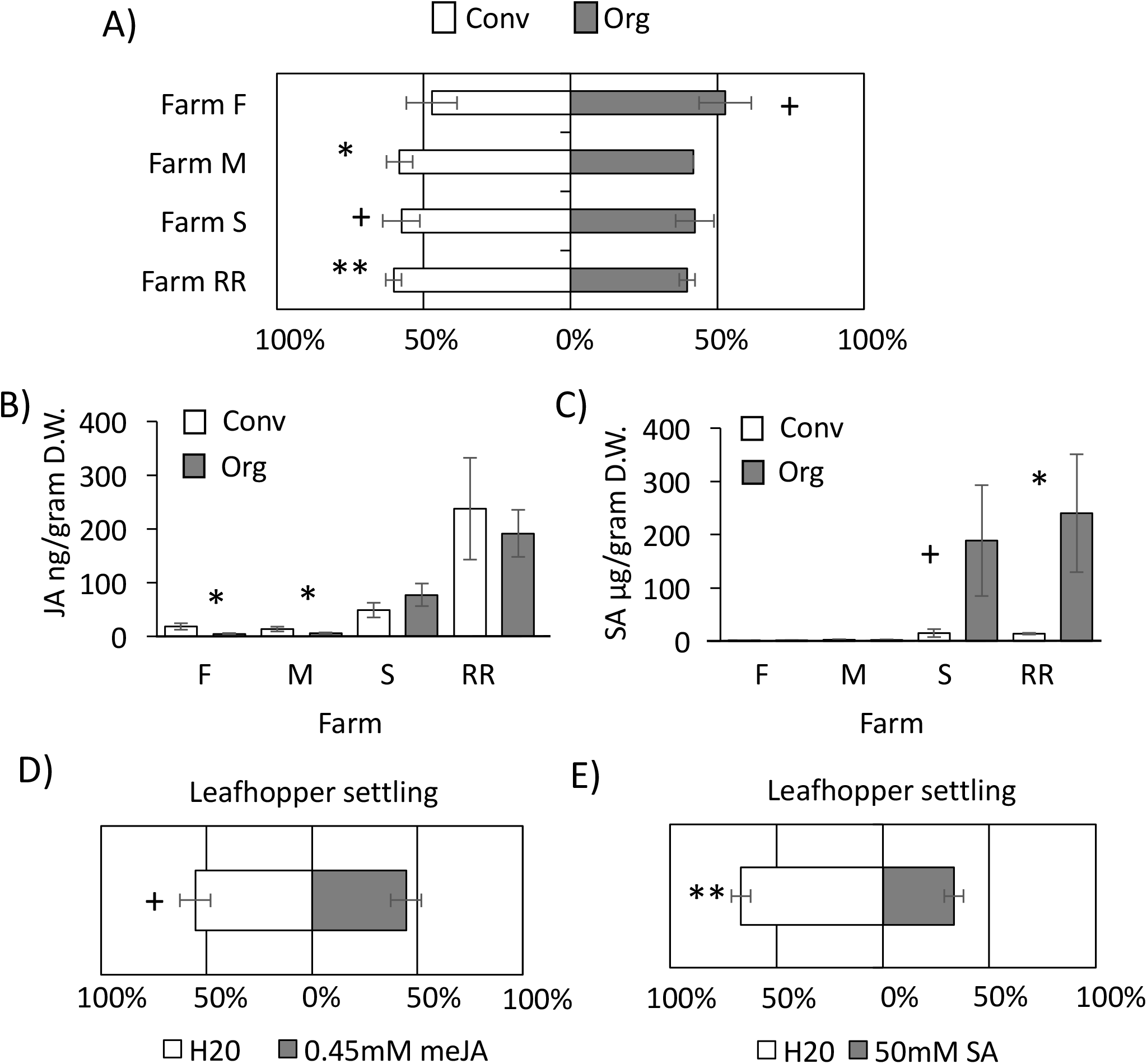
Organic management practices reduced insect settling and altered plant defense signaling pathways. (A) Leafhopper settling preference for leaves collected from Farm RR and three commercial processing tomato farms (Farm F, M, and S) in 2017. (B) Jasmonic acid (JA) and (C) salicylic acid (SA) content in tomato leaves from organic and conventional plots at Farm RR and three commercial processing tomato farms (Farm F, M, and S) in 2017. (D) Leafhopper settling on leaves induced with 0.45mM of methyl jasmonate (meJA) or with water as a control. (E) Leafhopper settling on leaves induced with 50mM of SA or with water as a control. (Mean ± SE; *N* = 18 for A, *N* = 6-12 for B and C, *N* = 15 for D and *N* = 12 for E). Binomial distribution test (A, D and E) and GLM at each site (B and C). Stars represent significant differences; +*P* < .1, \**P* < .05, \*\**P* < .001.

### Organic management practices altered plant defense signaling pathways

The phytohormones SA and jasmonic acid (JA) are important regulators of plant defense and changes often influence insect preference (28, 29). To determine if organic management practices may be altering SA or JA accumulation in tomato, we measured both phytohormones in leaves collected from all four sites. Although site-level variation was observed, leaves from organic farms had higher SA levels than those in conventional farms (*P* = 0.01275), driven by differences at Farm RR and S (Farm RR: ~17X more, *P* = 0.03495 and Farm S: ~12X more, *P* = 0.0629), but there were no differences between organic and conventional plots at Farm F and Farm M (Fig. 1C). No main effect of soil management on JA levels was observed (*P* = 0.49), but leaves from conventional plots on Farm F and M had elevated JA levels compared to the organic paired sites (Farm F: ~3X more, *P* = 0.0145 and Farm M: ~2X more, *P* = 0.0308) (Fig 1B). To determine if changes in SA or JA may be mediating leafhopper preference, we measured leafhopper settling on tomato leaves that had been induced with SA or methyl JA (meJA) compared to uninduced leaves in settling bioassays. Leafhoppers preferred to settle on control leaves compared to meJA- or SA-induced controls (Fig 1D; Fig 1E).

### Organic management practices altered plant and soil nutrient content

Organic and conventional management systems have drastically different soil fertility management. This can result in large variation in plant and soil nutrient content, which can directly or indirectly affect soil microbial populations and insect preference (30). We measured 14 different nutrients in leaves and soil collected at all four paired sites (Table S2, S3). There was considerable variation in plant nutrient content across the treatments and farms (Table S2, S3). Although nitrogen is one of the most limiting plant nutrients for insect herbivores, and often drives patterns of insect preference (31), there was no consistent difference in N content or C:N ratio in leaves between organic or conventional plots (Table S2). Sulfur and copper concentrations in leaves were higher in organically grown plants compared to conventional at three of the four sites (Table S2; Farm RR, Farm M, and Farm S). Conventionally managed soil had reduced total carbon, organic matter and sodium, and elevated magnesium, at three of the four sites compared to organically managed soils (Table S3).

### Rhizosphere microbial composition is associated with changes in plant nutrients and defense

Rhizosphere bacteria and fungi, which differ with management (32), have been previously shown to influence plant health by regulating defense compounds against insect herbivores (33). We examined if differences in rhizosphere communities were associated with observed differences in plant defense hormones in plants from the different farms. For both bacteria and fungi, tomato rhizosphere communities were more diverse under organic management at three of four sites (Fig 2A, B). Farms differed in microbial communities but organic and conventional communities remained distinct from each other at all sites (Fig 2C, D). Mantel tests were conducted to identify correlations among plant variables (nutrient, biomass, and hormone data) and microbes (bacteria or fungal composition). Plant variables were significantly associated with microbial community composition (RDA; Mantel Bacteria *r* = 0.33, *P* < .001; Fungi *r* = 0.51, *P* < .001). Because plant response variables were also associated with soil parameters (*P* < .001), we conducted a partial Mantel test to examine if microbial community composition remained significant after soil nutrients were included in the model. This analysis revealed that the structure of the microbial community, in particular the fungal community, was significantly associated with variation in plant traits including nutrient content and SA concentration, even when variation in soil nutrition was taken into account (partial Mantel *P* < .001).

**Fig. 2.**
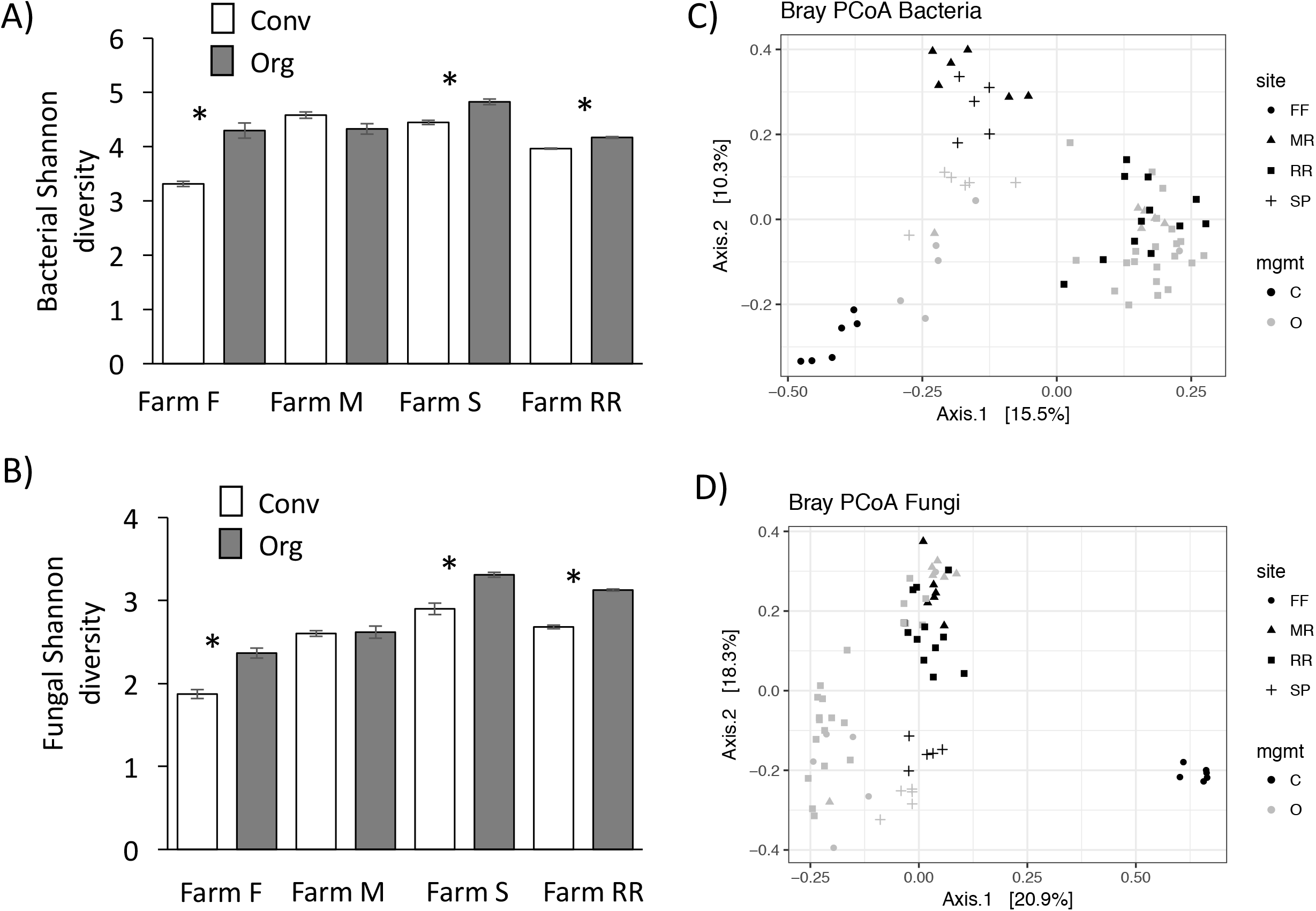
Bacterial and fungal diversity and community composition differ among organic and conventional sites. Rhizosphere microbial communities from processing tomato roots were sampled from paired organic (O) or conventional (C) farms at four locations and bacteria and fungi characterized using 16S and ITS metabarcoding. A) Bacterial and B) fungal diversity is greater in organic than conventional farms at three sites. C) Bacterial and D) fungal community composition differs among farms (PerMANOVA *P* < .001), between organic and conventional (*P* < .001) and according to the site: management interaction (*P* < .001). Mean ± SE; Stars represent significant differences; \**P* < .05).

To examine if any specific microbial amplicon sequence variants (ASVs) were associated with variation in plant SA concentrations, we performed a differential abundance analysis. ASVs from the bacterial genera *Bacillus*, *Ralstonia*, and *Exiguobacterium* were found in higher abundance when plants had high SA levels (Fig. 3; Table S4). This correlation suggests that variation in microbial communities is associated with variation in plant SA content.

**Fig. 3.**
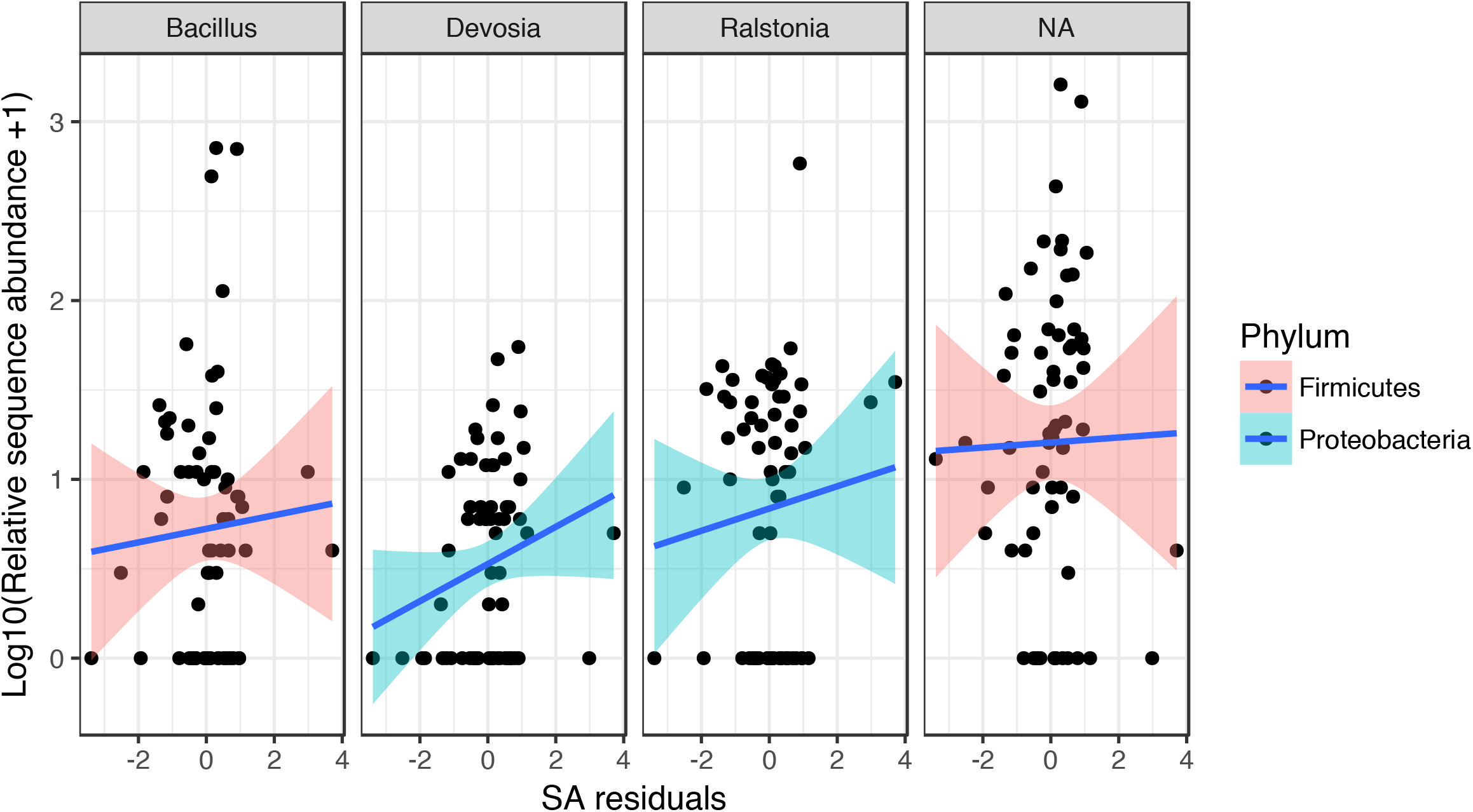
Microbial populations are associated with changes in salicylic acid. Relative abundance of root-associated bacterial taxa from 16S survey are positively related to the salicylic acid concentration in tomato leaves. Relationships are positive overall (*FDR* < 0.05) but driven by positive slope in conventional fields (top panels). Lines represent best-fit linear regression models. Bacterial genus names are across the top panel and “NA” indicates a taxon that could not be assigned to genus, but was a member of the Firmicutes.

### Soil biota drives differences in leafhopper preference and plant resistance

To isolate the relative importance of different soil components (physical structure, biological communities and chemical properties) in plant resistance, we performed a series of bioassays in the lab using rhizosphere soil collected from Russel Ranch, the farm where we observed the largest differences in insect populations, insect preference and plant resistance (Farm RR, Fig. 1, Fig. S1). Another reason we chose to focus on Russel Ranch for the soil slurries experiments was because it consists of 3 replicated plots for each management regime (34). To remove effects of soil physical properties, collected soils were washed and slurries from organic or conventional soils were used to inoculate tomato plants prior to bioassays. A greater number of leafhoppers settled on plants inoculated with conventional slurries compared to organic slurries (Fig. 4A), consistent with lab and field experiments (Fig. 1, Fig. S1). Leafhopper survival rate was higher on plants inoculated with slurries from the conventional plots, despite a sharp decline in survival over time in both treatments (Fig. 4B). These results suggest that management based regulation of plant resistance and insect preference may occur via soil biological or chemical parameters rather than physical properties at Russel Ranch.

**Fig. 4.**
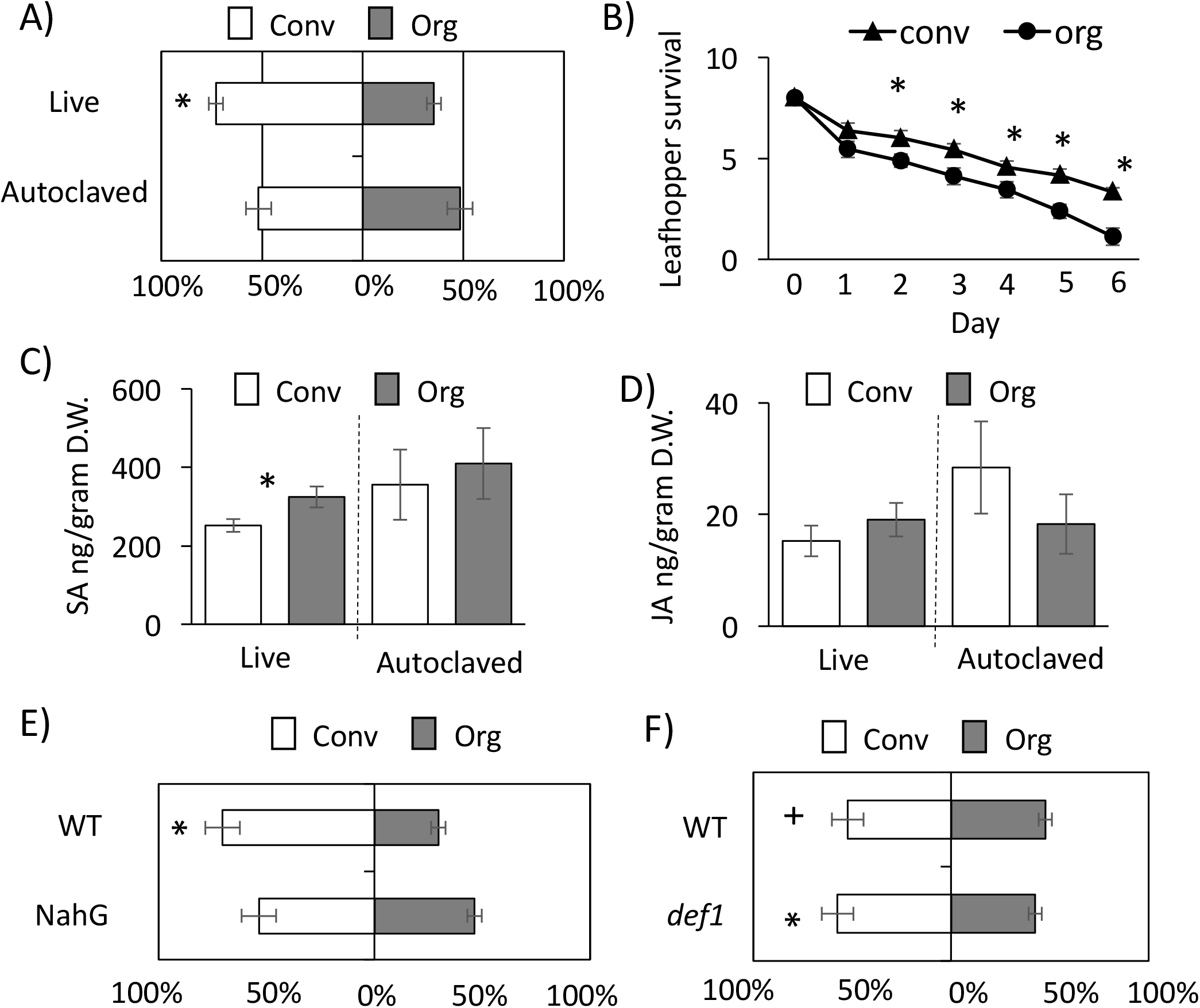
Soil biota drives differences in leafhopper populations, preference, and plant resistance. (A) Leafhopper settling preference for tomatoes grown a soil slurry prepared from conventional and organic rhizosphere soil from Russell Ranch that was untreated (Live) or autoclaved. (B) Leafhopper survival on tomatoes grown with a soil slurry prepared from conventional and organic rhizosphere soil from Farm RR. (C) Salicylic acid and (D) jasmonic acid content of leaves from tomatoes grown a soil slurry prepared from conventional and organic rhizosphere soil from Farm RR that was untreated or autoclaved. Leafhopper settling preference for leaves from (E) WT moneymaker or NahG and (F) WT castlemart or *def1* plants grown in conventional and organic soil slurries. Mean ± SE; *N* = 36 for A, *N* = 9 for B, *N* = 6-9 for C, D, E,F. Binomial distribution test (A, E and F), Student’s t-test (B, C and D). Stars represent significant differences; +*P* < .1, \**P* < .05.

Next, we investigated if soil biota within organically managed soils affect insect preference. Half of the slurry solution for each treatment (organic, conventional) was autoclaved to kill all microbes. When slurries were autoclaved (no live microbes), no difference in leafhopper settling preference was observed (Fig. 4A). These results suggest a critical role of soil microbes in mediating insect preference. Moreover, plants grown on the biologically active organic soil slurry had a 25% higher SA concentration compared to conventional, but no difference in the amount of SA was found when plants were inoculated with autoclaved slurries (Fig. 4C). Levels of JA did not differ between treatments (Fig. 4D). No significant differences in nutrient content between control and autoclaved soil slurries or organic and conventional treatments were observed (Table S5).

To determine if changes in SA or JA are responsible for differences in insect preference, we performed additional soil slurry experiments with NahG tomatoes, which are not able to accumulate SA and activate SA-mediated defenses, and *def1* tomatoes, in which JA signaling and related defenses are compromised (50). Leafhoppers had no preference between NahG plants grown using organic versus conventional soil slurries (Fig. 4E), while leafhoppers preferentially settled on *def1* and wt control plants that were grown in conventional soil slurries compared to the same plants grown in organic soil slurries (Fig. 4E, F). Differences were not as large for the *def1*/castlemart experiments (Fig. 4E), possibly due to cultivar differences or due to these experiments being conducted over a year after soil collection. These results collectively suggest that differences in microbial communities may mediate changes in insect preference and plant resistance levels through changes in SA signaling.

### Differences in soil properties drive changes in plant resistance across plant species

To determine if the impact of organic soil slurries on insect performance is conserved across plant species, we performed additional slurry experiments with *M. persicae*, a generalist hemipteran aphid and 3 additional plant species: carrot (*Daucus carota*), *Arabidopsis thaliana* and potato (*Solanum tuberosum*). Our results show that *M. persicae* fecundity is reduced on all three plants when grown with organic soil slurries compared to conventional (Fig. 5A). Lastly, we looked at the fitness of another type of herbivorous pest of tomato, *Manduca sexta*, which feeds by chewing, as opposed to phloem feeding, as hemipterans do. There was no significant difference in dry weight of *M. sexta* between treatments (Fig. 5B).

**Fig. 5.**
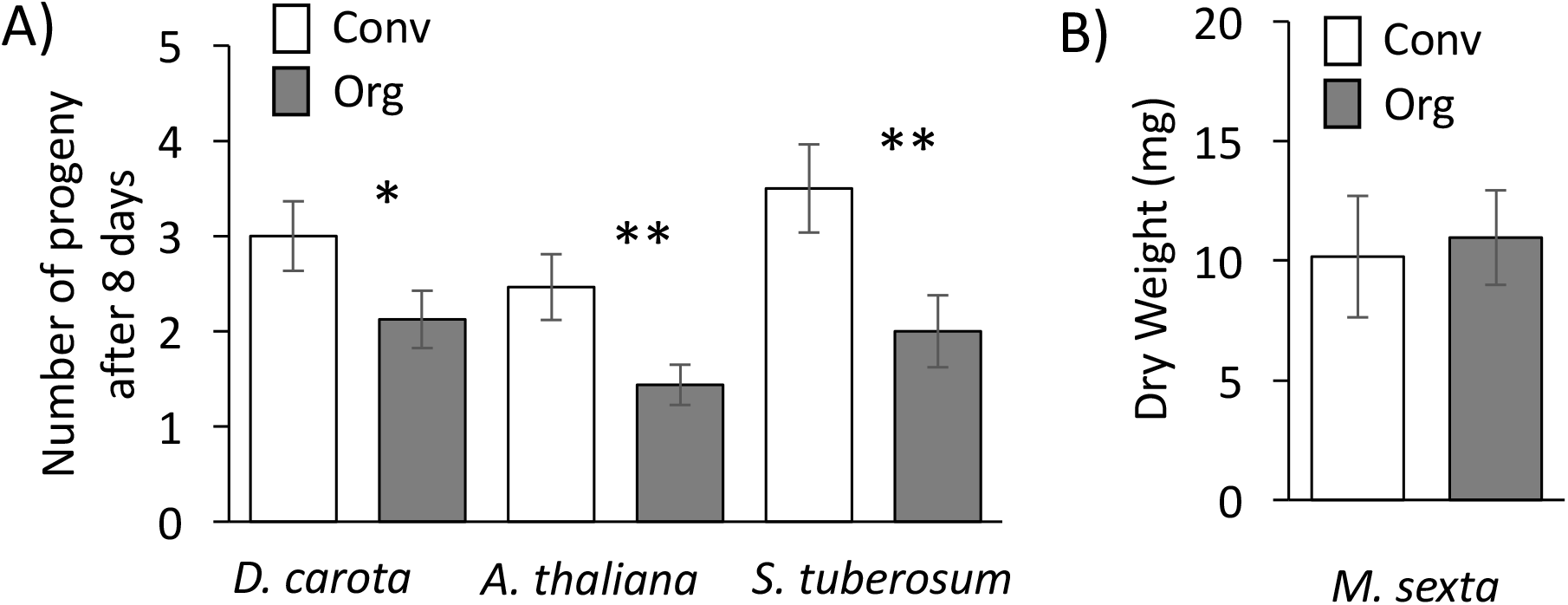
Soil biota drive differences in hemipteran population growth across plant species. (A) *Myzus persicae* fecundity on carrot (*Daucus carota*), Arabidopsis (*A. thaliana*), and potato (*Solanum tuberosum*) inoculated with soil slurries prepared from conventional and organic rhizosphere soil at Farm RR. (C) *Manduca sexta* dry weight when reared on tomatoes inoculated with soil slurries prepared from conventional and organic plots at RR. Mean ± SE; *N* = 12-18 for A and *N* = 9-13 for B. Student’s t-test (A-B). Stars represent significant differences; \**P* < .05, \*\**P* < .01, \*\*\**P* < .001.

## Discussion

Our results demonstrate that tomatoes grown on long-term organic farms have lower insect populations (Fig. S1, B) and that fewer leafhopper pests settle on these plants (Fig. 1A) compared to conventionally-grown plants. We show that organic soil management promoted SA accumulation, which directly influences plant-insect interactions (Fig. 1B-E). We demonstrate that changes in SA and insect preference are dependent on shifts in soil microbial communities associated with long-term organic management (Fig. 1, 2, 3, 4), and that these findings may be applicable to multiple plant systems (Fig. 5). Although soil microbial effects on plant pathogens and soil-borne pests in agroecosystems are appreciated and relatively well-described, here we show that soil microbial communities likely play an unappreciated role in depressing insect pest populations through changes in plant resistance. These results suggest that more sustainable insect management strategies can be developed through soil health management.

Plants have evolved complex immune systems to protect themselves against pests and pathogens. Previous studies have identified SA in mediating plant defense responses to hemipterans (36–38), while changes in JA and ethylene (ET) have been largely connected to defenses against chewing insects (28). Consistent with this work, we observed an impact of organic soil management on plant resistance to multiple hemipterans (Fig. 1, 4, 5), and no impact on the chewing caterpillar *M. sexta* (Fig. 5), when SA levels were elevated. Despite JA and SA’s induction of alternative resistance pathways, there is evidence to suggest that considerable crosstalk exists and that both can contribute to resistance against the same attacker. For example, aphids were shown to induce the JA pathway in addition to the SA pathway and to also be susceptible to JA-mediated plant defenses (39, 40). We observed that treating plants with meJA repelled leafhoppers in lab bioassays (Fig. 1D), though there was no significant difference in JA levels in plants from Russel Ranch (Fig. 1B) or in plants inoculated with biologically active or inert slurries (Fig. 4D). Furthermore, leafhoppers were still repelled from jasmonate-deficient plants grown in organic soil slurries as compared to slurries from conventionally managed soil (Fig. 4F). Together, these results suggest that changes in SA are primarily driving changes in leafhopper-plant interactions in our system.

Despite knowledge of the essential roles that microbe communities play in agroecosystems, we still have a limited understanding of the direct benefits that microbial diversity and composition provide in terms of plant health and resistance to insect pests. Organic sites in our study exhibited an overrepresentation of specific microbial taxa which are known to be involved in the induction of plant defenses (41, 42), including *Pseudomonas*, *Ochrobactrum*, *Glutamicibacter, Bacillus, Ralstonia*, and others (Table S4). Furthermore, microbial taxa from the bacterial genera *Bacillus* and *Ralstonia* were associated with variation in plant SA concentrations in our field experiments (Fig. 3). The presence of these particular taxa may promote plant-induced resistance (43, 44) though plant induction of SA, or changes in the soil environment may also modulate microbial interactions directly (45). In our study, organically managed soils had higher organic matter and carbon, and reduced magnesium, compared to paired conventionally managed soils at three of the four sites (Table S3). Changes in soil chemistry or nutrient availability in organic soils may contribute to enhanced plant defense responses through changes in the soil microbiome (46). Although the particular microbial taxa or community composition underlying this effect are currently unknown, this study strongly suggests that organic practices in agro-ecosystems can promote plant resistance to insect pests through changes in soil microbiota.

While it is known that soil microbes can influence above-ground plant-insect interactions through changes in plant signaling and defense (47-50), the management techniques that promote beneficial microbial populations remain poorly understood. Our data demonstrate that organic management practices alter soil microbial communities, alter plant defense potentials through changes in SA, and influence hemipteran settling and performance (Fig. 1, 2, 3). Although we cannot distinguish effects of diversity per se vs. compositional changes or specific taxa underlying this effect, laboratory assays strongly implicate soil microbiota in plant protection (Fig. 4). Field surveys support the hypothesis that organic practices can influence insect preference at large scales, but also suggest that variation in practices or local conditions may moderate these results in some locations (Fig. 1, 2, Fig. S1). Although further work is required to dissect the particular mechanisms involved, including investigation of microbial strains or signals, our results suggest that healthy soils cultivated using organic practices can promote sustainable and resilient yields in the face of hemipteran pest pressure. Organic agriculture therefore holds great potential to broadly improve the delivery of key ecosystem services critical for the sustainability of farming systems and the resilience of the food supply.

## Acknowledgements

The authors would like to thank Franz Bender, Jesper Richardy, Griffin Hall, Samuel Tookey, UCCE farm advisors, Russell Ranch staff and growers for participating in this study, assisting with sampling. This work was supported by start-up funds from UC Davis to CC, AG, and RV; the California Tomato Research Institute to AG, CC and RV, the California Potato Research Advisory Board to CC, and the USDA-NIFA, Agricultural Experiment Station Project #CA-D-PLS-2332-H to AG.

## Author contributions

R.B., A.L.C., J.E.S, and A.I. conducted most of the experiments and analysis. C.L.C., A.G. and R.L.V. designed all experiments and directed the project. R.B. C.L.C., A.G. and R.L.V. wrote the paper with comments and input from all authors.

---

**Supplemental Fig. 1.**
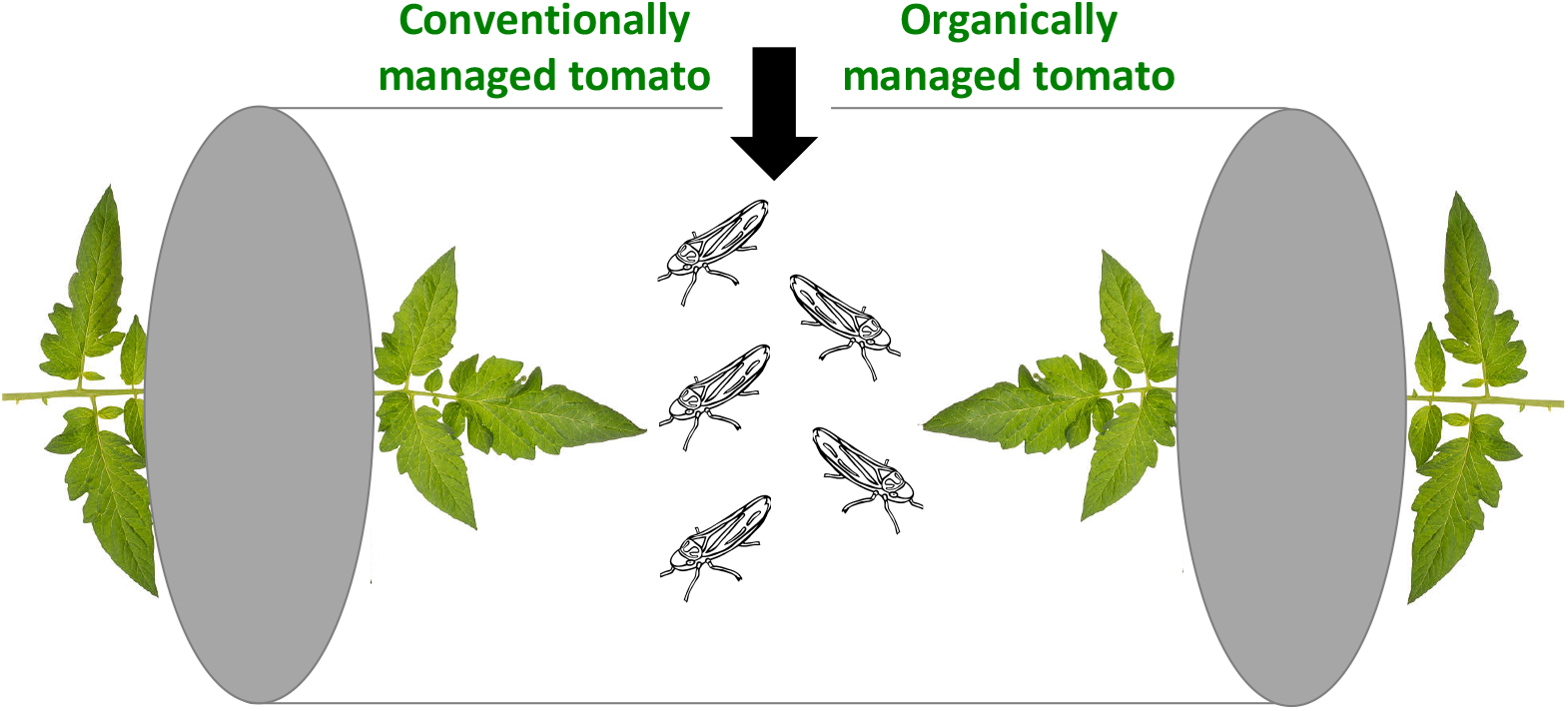
Diagrammatic representation of choice bioassay. At one end of a clear cylinder, a conventionally managed tomato leaf is sealed inside using foam whilst an organically managed tomato leaf is sealed inside the other end. Five Avirulent beet leafhoppers are starved 2 hours prior to the experiment and inserted into the center of the cylinder.

**Supplemental Fig. 2.**
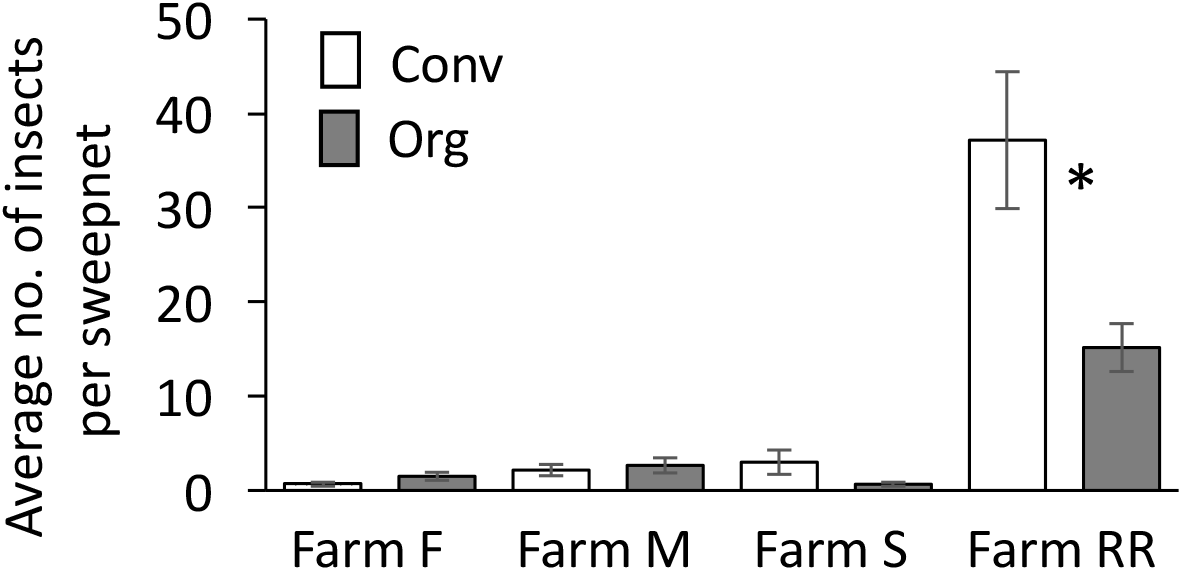
Organic management practices reduced insect populations in processing tomatoes. The number of insects collected in sweepnets on organic and conventional plots at Farm RR and three commercial processing tomato farms (Farm F, M, and S) in 2017. Mean ± SE; *N* = 6. Whitney U test. Stars represent significant differences; \**P* < .05).

**Supplemental Table 1.**
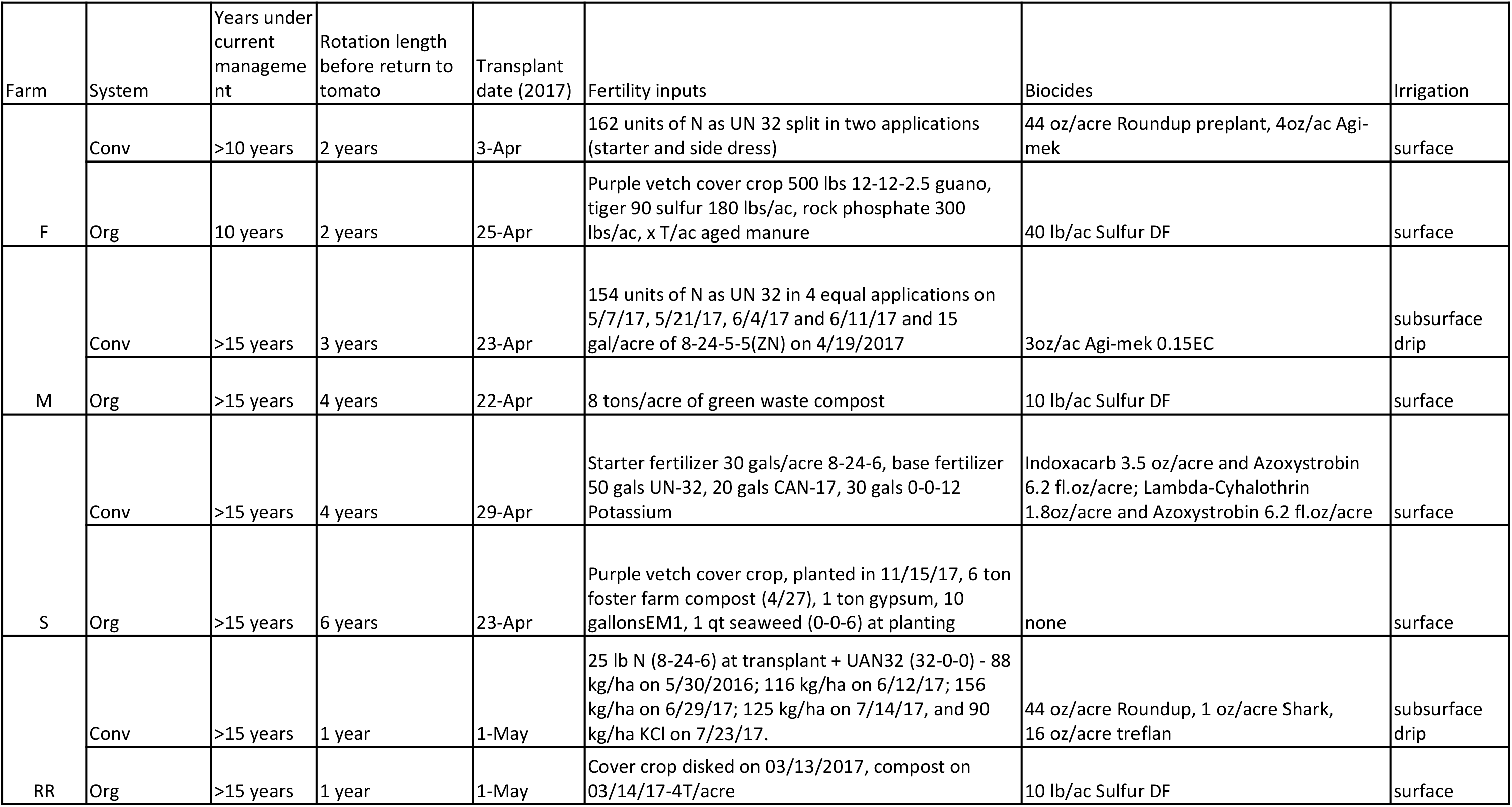
Field management

| Farm | System | Years under current management | Rotation length before return to tomato | Transplant date (2017) | Fertility inputs | Biocides | Irrigation |
| --- | --- | --- | --- | --- | --- | --- | --- |
| F | Conv | >10 years | 2 years | 3-Apr | 162 units of N as UN 32 split in two applications (starter and side dress) | 44 oz/acre Roundup preplant, 4oz/ac Agi-mek | surface |
|  | Org | 10 years | 2 years | 25-Apr | Purple vetch cover crop 500 lbs 12-12-2.5 guano, tiger 90 sulfur 180 lbs/ac, rock phosphate 300 lbs/ac, x T/ac aged manure | 40 lb/ac Sulfur DF | surface |
| M | Conv | >15 years | 3 years | 23-Apr | 154 units of N as UN 32 in 4 equal applications on 5/7/17, 5/21/17, 6/4/17 and 6/11/17 and 15 gal/acre of 8-24-5-5(ZN) on 4/19/2017 | 3oz/ac Agi-mek 0.15EC | subsurface drip |
|  | Org | >15 years | 4 years | 22-Apr | 8 tons/acre of green waste compost | 10 lb/ac Sulfur DF | surface |
| S | Conv | >15 years | 4 years | 29-Apr | Starter fertilizer 30 gals/acre 8-24-6, base fertilizer 50 gals UN-32, 20 gals CAN-17, 30 gals 0-0-12 Potassium | Indoxacarb 3.5 oz/acre and Azoxystrobin 6.2 fl.oz/acre; Lambda-Cyhalothrin 1.8oz/acre and Azoxystrobin 6.2 fl.oz/acre | surface |
|  | Org | >15 years | 6 years | 23-Apr | Purple vetch cover crop, planted in 11/15/17, 6 ton foster farm compost (4/27), 1 ton gypsum, 10 gallons EM1, 1 qt seaweed (0-0-6) at planting | none | surface |
| RR | Conv | >15 years | 1 year | 1-May | 25 lb N (8-24-6) at transplant + UAN32 (32-0-0) - 88 kg/ha on 5/30/2016; 116 kg/ha on 6/12/17; 156 kg/ha on 6/29/17; 125 kg/ha on 7/14/17, and 90 kg/ha KCl on 7/23/17. | 44 oz/acre Roundup, 1 oz/acre Shark, 16 oz/acre treflan | subsurface drip |
|  | Org | >15 years | 1 year | 1-May | Cover crop disked on 03/13/2017, compost on 03/14/17-4T/acre | 10 lb/ac Sulfur DF | surface |

**Supplemental Table 2.** Plant nutrient content of tomato leaves grown under organic and conventional management

|  | Farm F |  | Farm M |  | Farm S |  | Farm RR |  |
| --- | --- | --- | --- | --- | --- | --- | --- | --- |
|  | Conv | Org | Conv | Org | Conv | Org | Conv | Org |
| **Shoot dry weight** | 6.97*** | 4.18 | 42.61 | 45.64 | 16.56 | 18.52 | 47.92* | 56.05 |
| **C%** | 36.98*** | 39.23 | 38.91** | 37.33 | 30.44 | 32.56 | 39.14** | 38.65 |
| **N%** | 4.75*** | 5.42 | 5.26*** | 3.81 | 4.07 | 4.47 | 4.74* | 4.87 |
| **P%** | 0.51 | 0.48 | 0.49 | 0.45 | 0.31 | 0.34 | 0.54* | 0.57 |
| **K%** | 3.06 | 2.77 | 2.66* | 2.26 | 1.21* | 1.51 | 2.02*** | 2.41 |
| **Ca%** | 2.67*** | 3.58 | 3.58 | 3.58 | 2.21* | 2.98 | 2.93*** | 3.44 |
| **Mg%** | 1.32*** | 1.47 | 1.24 | 1.31 | 0.96* | 0.88 | 1.99*** | 1.89 |
| **S%** | 0.42*** | 0.98 | 0.87*** | 1.28 | 0.35*** | 1.00 | 0.71*** | 1.27 |
| **Mn mg/kg** | 142.08* | 121.08 | 96.17 | 91.67 | 140.67 | 142.92 | 183.25*** | 118.22 |
| **Fe mg/kg** | 3579.17*** | 1055.92 | 1571.67 | 2076.75 | 6677.33 | 5332.75 | 1415.08*** | 1000.50 |
| **Cu mg/kg** | 17.00 | 17.00 | 24.00*** | 30.58 | 26.25** | 31.00 | 18.86*** | 24.19 |
| **B mg/kg** | 102.67*** | 75.42 | 201.58* | 164.25 | 45.33 | 46.50 | 78.86** | 75.03 |
| **Al mg/kg** | 2151.00*** | 586.83 | 847.67 | 1127.58 | 4817.83 | 3869.25 | 719.06*** | 487.86 |
| **Zn mg/kg** | 27.00*** | 33.75 | 26.00 | 26.67 | 53.83*** | 43.08 | 30.58 | 30.58 |
| **Na mg/kg** | 631.08** | 486.75 | 307.67* | 250.83 | 429.67 | 357.00 | 493.58 | 535.58 |

Mean, $N = 12$ ; Stars represent significant differences of nutrients between conventional and organic paired sites, \* $P < .1$ , \*\* $P < .05$ , \*\*\* $P < .001$ . Red and blue cell shading represent significant increases or decreases in the conventional system relative to organic system respectively.

**Supplemental Table 3.**
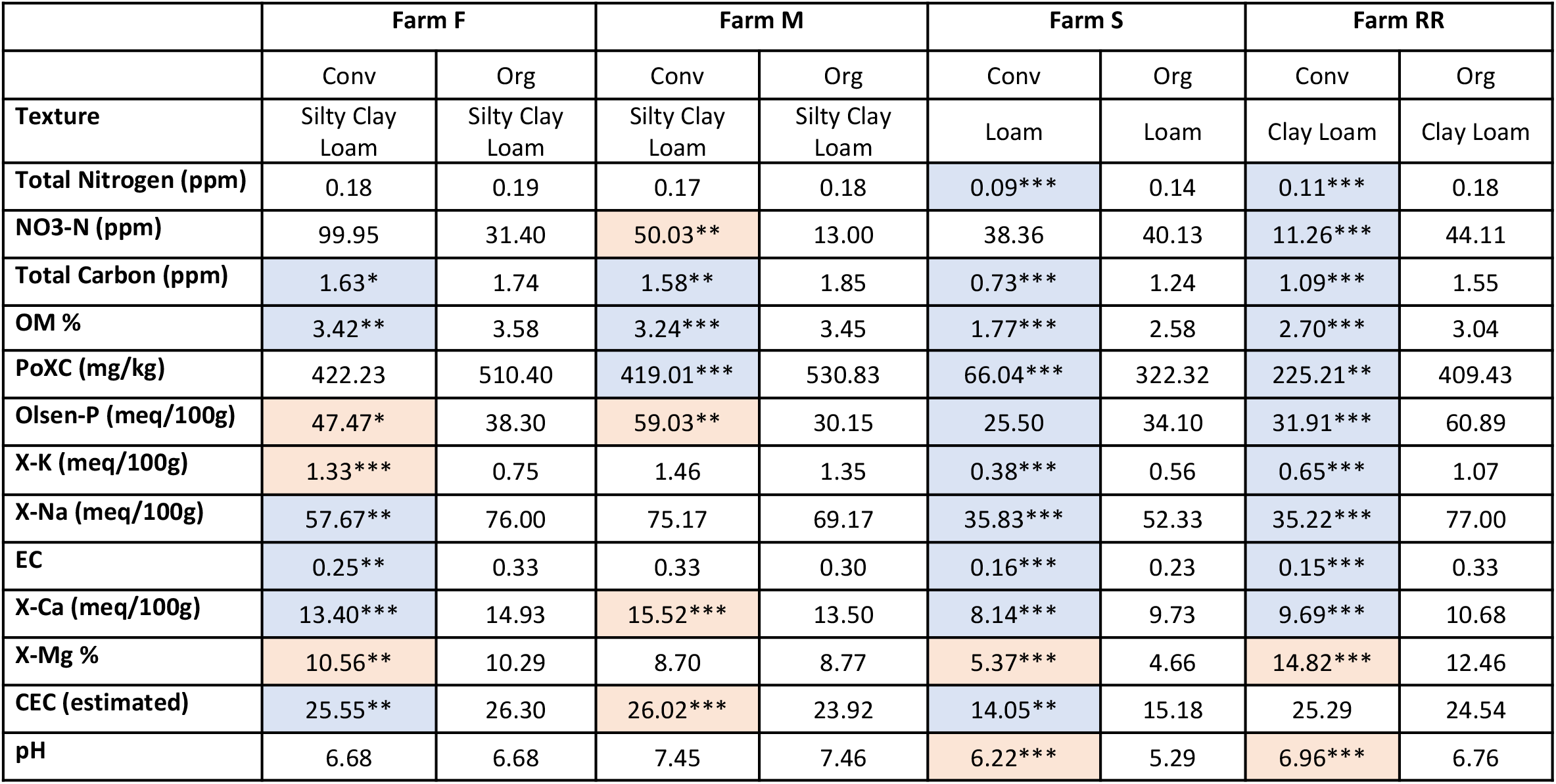
Organic management influences soil fertility and regulators of microbial communities. Means are shown (*N* = 6 for farm F, M and S; *N* =18 for RR) Stars represent significant differences between organic and conventional soils for each farm, \**P* < .05, \*\**P* < .01. *** *P* ≤ .001. Red and blue cell shading represent signifiant increases or decreases in the conventional system relative to organic system respectively. Ppm = parts per millions, meq/100g= millequivalents per 100 grams of soil. OM = Organic matter, PoxC = Permanganate Oxidizable Carbon, CEC = cation exchange capacity, EC= Electrical Conductivity.

|  | Farm F |  | Farm M |  | Farm S |  | Farm RR |  |
| --- | --- | --- | --- | --- | --- | --- | --- | --- |
|  | Conv | Org | Conv | Org | Conv | Org | Conv | Org |
| **Texture** | Silty Clay Loam | Silty Clay Loam | Silty Clay Loam | Silty Clay Loam | Loam | Loam | Clay Loam | Clay Loam |
| **Total Nitrogen (ppm)** | 0.18 | 0.19 | 0.17 | 0.18 | 0.09*** | 0.14 | 0.11*** | 0.18 |
| **NO3-N (ppm)** | 99.95 | 31.40 | 50.03** | 13.00 | 38.36 | 40.13 | 11.26*** | 44.11 |
| **Total Carbon (ppm)** | 1.63* | 1.74 | 1.58** | 1.85 | 0.73*** | 1.24 | 1.09*** | 1.55 |
| **OM %** | 3.42** | 3.58 | 3.24*** | 3.45 | 1.77*** | 2.58 | 2.70*** | 3.04 |
| **PoXC (mg/kg)** | 422.23 | 510.40 | 419.01*** | 530.83 | 66.04*** | 322.32 | 225.21** | 409.43 |
| **Olsen-P (meq/100g)** | 47.47* | 38.30 | 59.03** | 30.15 | 25.50 | 34.10 | 31.91*** | 60.89 |
| **X-K (meq/100g)** | 1.33*** | 0.75 | 1.46 | 1.35 | 0.38*** | 0.56 | 0.65*** | 1.07 |
| **X-Na (meq/100g)** | 57.67** | 76.00 | 75.17 | 69.17 | 35.83*** | 52.33 | 35.22*** | 77.00 |
| **EC** | 0.25** | 0.33 | 0.33 | 0.30 | 0.16*** | 0.23 | 0.15*** | 0.33 |
| **X-Ca (meq/100g)** | 13.40*** | 14.93 | 15.52*** | 13.50 | 8.14*** | 9.73 | 9.69*** | 10.68 |
| **X-Mg %** | 10.56** | 10.29 | 8.70 | 8.77 | 5.37*** | 4.66 | 14.82*** | 12.46 |
| **CEC (estimated)** | 25.55** | 26.30 | 26.02*** | 23.92 | 14.05** | 15.18 | 25.29 | 24.54 |
| **pH** | 6.68 | 6.68 | 7.45 | 7.46 | 6.22*** | 5.29 | 6.96*** | 6.76 |

**Supplementary Table 4.** Bacterial taxa differentially abundant on tomato roots between fields in organic vs conventional soil management. Taxa were identified using DESeq2. Taxa with a log2fold Change greater than 1 were more abundant in Organic fields, whereas those less than one were more abundant in conventional fields.

| Kingdom | Phylum | Class | Order | Family | Genus | SequenceID | baseMean | Log2 FoldChange | False Discovery Rate |
| --- | --- | --- | --- | --- | --- | --- | --- | --- | --- |
| Bacteria | Proteobacteria | Gammaproteobacteria | Xanthomonadales | Xanthomonadaceae | Rhodanobacter | Seq87 | 25.95159286 | -5.552186069 | 6.71E-06 |
| Bacteria | Proteobacteria | Betaproteobacteria | Burkholderiales | Oxalobacteraceae | Janthinobacterium | Seq23 | 43.60225996 | -5.287749644 | 0.043433299 |
| Bacteria | Proteobacteria | Gammaproteobacteria | Enterobacteriales | Enterobacteriaceae | Yersinia | Seq14 | 336.199005 | -5.028759101 | 3.55E-06 |
| Bacteria | Proteobacteria | Gammaproteobacteria | Xanthomonadales | Xanthomonadaceae | Rhodanobacter | Seq55 | 112.5465685 | -5.020068278 | 0.043433299 |
| Bacteria | Bacteroidetes | Flavobacteriia | Flavobacteriales | Flavobacteriaceae | Flavobacterium | Seq73 | 11.7358541 | -4.245670914 | 0.03884778 |
| Bacteria | Proteobacteria | Gammaproteobacteria | Pseudomonadales | Pseudomonadaceae | Pseudomonas | Seq7 | 1833.758731 | -3.15640054 | 6.71E-06 |
| Bacteria | Proteobacteria | Alphaproteobacteria | Sphingomonadales | Sphingomonadaceae | Sphingobium | Seq176 | 7.641398516 | -2.969971736 | 0.015174083 |
| Bacteria | Proteobacteria | Gammaproteobacteria | Enterobacteriales | Enterobacteriaceae | NA | Seq5 | 1904.823229 | -2.557292633 | 0.005174835 |
| Bacteria | Actinobacteria | Actinobacteria | Corynebacteriales | Nocardiaceae | Rhodococcus | Seq48 | 19.40098497 | 1.894006416 | 5.32E-05 |
| Bacteria | Firmicutes | Bacilli | Bacillales | Paenibacillaceae | Paenibacillus | Seq16 | 287.1503902 | 1.941977892 | 0.000388882 |
| Bacteria | Firmicutes | Bacilli | Bacillales | Family_XII | Exiguobacterium | Seq75 | 11.98388315 | 1.982208331 | 0.033586704 |
| Bacteria | Proteobacteria | Alphaproteobacteria | Rhizobiales | Rhizobiaceae | NA | Seq33 | 51.51773028 | 2.033100265 | 0.007010338 |
| Bacteria | Actinobacteria | Actinobacteria | Micrococcales | Sanguibacteraceae | Sanguibacter | Seq101 | 8.100463395 | 2.22622812 | 0.034840877 |
| Bacteria | Actinobacteria | Actinobacteria | Micrococcales | Microbacteriaceae | Microbacterium | Seq41 | 36.38781913 | 2.2512469 | 1.76E-08 |
| Bacteria | Proteobacteria | Alphaproteobacteria | Rhizobiales | Methylobacteriaceae | Microvirga | Seq103 | 13.85211138 | 2.557406515 | 0.00331975 |
| Bacteria | Proteobacteria | Gammaproteobacteria | Pseudomonadales | Pseudomonadaceae | Pseudomonas | Seq29 | 54.10603478 | 2.599161574 | 0.015174083 |
| Bacteria | Firmicutes | Bacilli | Bacillales | Bacillaceae | Bacillus | Seq49 | 19.29518115 | 3.014437507 | 0.000748747 |
| Bacteria | Bacteroidetes | Sphingobacteriia | Sphingobacteriales | Sphingobacteriaceae | Pedobacter | Seq32 | 65.22587825 | 3.65153209 | 0.00331975 |
| Bacteria | Firmicutes | Bacilli | Bacillales | Bacillaceae | NA | Seq60 | 33.17256696 | 3.787099683 | 6.71E-06 |
| Bacteria | Proteobacteria | Alphaproteobacteria | Rhizobiales | Bradyrhizobiaceae | Bosea | Seq449 | 3.204729918 | 3.894881668 | 0.049766386 |
| Bacteria | Proteobacteria | Gammaproteobacteria | Pseudomonadales | Pseudomonadaceae | Pseudomonas | Seq131 | 3.08617392 | 4.065776492 | 0.025730032 |
| Bacteria | Proteobacteria | Gammaproteobacteria | Xanthomonadales | Xanthomonadaceae | Stenotrophomonas | Seq68 | 5.705639918 | 4.502285298 | 0.004464205 |
| Bacteria | Actinobacteria | Actinobacteria | Micrococcales | Microbacteriaceae | Plantibacter | Seq124 | 4.268012173 | 4.709319603 | 5.26E-10 |
| Bacteria | Proteobacteria | Alphaproteobacteria | Rhizobiales | Rhizobiaceae | Kaistia | Seq138 | 5.569611401 | 4.716490074 | 0.001083245 |

**Supplemental Table 5.** Soil slurry composition

| System | Treatment | pH<br>(mg/L) | NH4-N<br>(mg/L) | NO3-N<br>(mg/L) | K<br>(Total)<br>(mg/L) | Ca<br>(Total)<br>(mg/L) | Mg<br>(Total)<br>(mg/L) | Na<br>(Total)<br>(mg/L) | Zn<br>(Total)<br>(mg/L) | Mn<br>(Total)<br>(mg/L) | Fe<br>(Total)<br>(mg/L) | Cu<br>(Total)<br>(mg/L) |
| --- | --- | --- | --- | --- | --- | --- | --- | --- | --- | --- | --- | --- |
| Org | Autoclaved | 6.97 | 2.17 | 93.87 | 44.73 | 59.10 | 82.57 | 12.90 | 0.14 | 1.23 | 55.06 | 0.06 |
|  | Control | 6.67 | 2.32 | 87.67 | 39.87 | 52.07 | 62.27 | 10.17 | 0.00 | 0.34 | 9.92 | 0.00 |
| Conv | Autoclaved | 7.08 | 2.68 | 88.77 | 45.70 | 59.43 | 62.67 | 14.90 | 0.03 | 0.49 | 19.01 | 0.02 |
|  | Control | 6.89 | 2.68 | 83.83 | 45.90 | 59.48 | 64.87 | 18.42 | 0.05 | 0.63 | 28.02 | 0.04 |

## Materials and Methods

### Field study sites

Field studies took place during the 2017 growing season at the organic and conventional long-term treatments of the Century experiment established in 1993 at Russell Ranch (Davis, CA, USA) (34). Three additional field studies took place on commercial farms in Yolo county in 2017. In these studies, paired long-term organic and conventional processing tomato field plots were compared. At Russel Ranch, paired sites refer to both the organic and conventional replicated plots on a research farm (34). For the other farms, paired sites refer to organic and conventional fields being managed by the same grower in nearby plots, and where tomato was sown at the same time. Details of field management strategies are available in Table S1. At Russell Ranch each treatment plot was 0.4 ha and replicated three times in completely randomized block design. For commercial farms twelve sampling locations per field were selected randomly for sampling. Details on soil chemistry for each site are available in Table S3.

### Insect sweepnet sampling

Insect populations were sampled at the study sites described above three weeks after transplanting. To standardize collections, six areas of each plot were collected from along a transect. Each collection consisted of ten sweeps up and down the field within an eight-row boundary along the transect. Samples were bagged and frozen until insects were counted and sorted to order.

### Plants and growth conditions

Moneymaker cv. tomato, castlemart cv. tomato, transgenic NahG tomato in the moneymaker background (35), and the jasmonate-deficient *def1* mutant tomato in the castlemart background (51) were used in lab studies, while Heinz 8504 cv. tomato was used for all Russell Ranch experiments. For commercial farms tomato cultivar varied by site (See Table S1). For *Arabidopsis*, potato and carrot experiments, Col-0, Desiree and Sativus cultivars were used respectively. For controlled experiments, plants were grown in Conviron growth chambers under 25°C/20°C day/night with a photoperiod of 16/8 hr day/night at a relative humidity of 50% and a light intensity of 200 mmol m^−2^ sec^−1^. The same growth conditions were used in all subsequent experiments.

### Insects

Avirulent beet leafhoppers, *Circulifer tenellus* were reared on beet (*Beta vulgaris*) under controlled conditions (28°C/24°C day/night with a photoperiod of 16/8 hr day/night). Aphids (*Myzus persicae*) were reared on potato under controlled conditions (28°C/24°C day/night with a photoperiod of 16/8 hr day/night). *Manduca sexta* eggs were ordered from Carolina Biological Supply (Burlington, N.C.) and held at room temperature until hatching. Neonates were immediately used in bioassays under controlled conditions and not reared.

### Soil slurries experiments

Rhizosphere soil was collected separately from the three-replicated conventional and organic plots at Russell Ranch Century Experiment. We chose to focus on Russel Ranch as a source of soils for our slurry experiment because 1) results were most contrasting at this site while being representative of other farms and 2) it consists of 3 replicated plots for each management regime with consistent management for the last 25 years. Soil slurries were prepared by mixing sampled soil with ¼ strength Hoagland’s nutrient solution at 1g soil to 5ml solution for one hour at 350 rpms. The solution was then left to settle for 1 hour at room temperature and centrifuged at 500 rpm for 5 mins. After centrifuging, the supernatant was removed and either autoclaved at 120°C for 30 minutes or left untreated. The soil slurries were added at the time of sowing at a rate of 15ml twice per week until the settling bioassay, fecundity of survival experiment was performed three weeks after seedling emergence. This methodology was used for all plant species.

### Settling bioassays from the field

Tomato branches were collected from the conventional and organic locations mentioned above three weeks after transplanting and immediately used for settling bioassays. Avirulent beet leafhoppers were collected and starved for two hours prior to the experiment. A cage was constructed that allowed an organic tomato leaf to be sealed at one end and a conventional leaf in the other (Fig. S1). Developmentally similar leaves were selected to standardize the assay. Five avirulent beet leafhoppers were introduced in the center of the cage equidistant to both leaves. Leafhopper position was recorded two hours after release. This time was chosen based on preliminary experiments where leafhopper settling was measured at multiple time points over 24 hours. The settling bioassays were conducted in the dark so leafhoppers could not make a settling based on visual cues. Each experiment was repeated 18 times. See Fig. S1 for experimental design of the settling experiment.

### Settling bioassays with hormone induced plants

Tomatoes were treated three weeks after tomato emergence. For salicylic acid induction 2 grams of salicylic acid was dissolved in 250 ml of H_2_O containing 0.1% Tween 20 and sprayed on plants until run off. For jasmonic acid induction, a solution of methyl jasmonate 95% (0.45 mM methyl JA with 0.1% Tween 20) was used. Control plants were treated with H_2_O containing 0.1% Tween 20. All settling bioassays were conducted as described above, 24 hours after chemical treatment. Each experiment was repeated 12-18 times. See Fig. S1 for experimental design of the settling experiment.

### Settling bioassays with soil slurries

Tomatoes were grown in sterile soil supplemented with soil slurries as described above. Three weeks after tomato emergence, settling bioassays were performed as described above. Each experiment was repeated 36 times. See Fig. S1 for experimental design of the settling experiment.

### Survival bioassays and fecundity assays

Tomatoes were grown in sterile soil supplemented with soil slurries as described above. Three weeks after tomato emergence, 8 adult *C. tenellus* were installed on a single leaf and survival was recorded daily over 6 days. Each experiment was repeated at least 2 times. *D. carota, A. thaliana* and *S. tuberosum* were grown in sterile soil supplemented with soil slurries as described above. At three weeks post-emergence, one adult *M. persicae* aphid was placed on a leaf. After 24 hours, all aphids except one nymph were removed. After 9 days, the progeny of the founder nymph, which was now an adult, were counted to determine fecundity. Each experiment contained at least 6 replicates and was repeated at least 2 times.

### *Manduca sexta* weight gain

After *M. sexta* emerged from eggs, neonate larva were immediately moved to cages with a paintbrush. Cages were installed on three-week-old tomatoes, post-emergence, that were grown in sterile soil supplemented with soil slurry twice as described above. One week later all caterpillars were removed, freeze dried, and weighed. Each experiment was repeated at least 2 times with at least 9 replicates.

### Phytohormone extraction and LC-MS analysis

During sweepnet sampling, developmentally similar true leaves from 6 separate three-week-old tomato plants in each plot were removed and immediately frozen in liquid nitrogen. For the plants grown in the soil slurries, developmentally similar true leaves were removed from tomato leaves three weeks post-emergence and immediately frozen in liquid nitrogen. Samples were stored at −80°C until they were lyophilized. Subsequent tissue was then weighed, ground in a Harbil paintshaker (Fluidman, Wheeling, IL) and extracted according to Casteel et al 2016 (52). Samples were run on an Agilent 6420A Triple-quadrupole MS with an Infinity II HPLC (Agilent Technologies, Santa Clara, CA). Quantification was based on an isotopically labelled internal standard that was spiked in each sample before the extraction. At least 9 samples were measured for each treatment. For phytohormone quantification no insects were used, which means our data represent “constitutive levels”, however, all field samples have some level of damage.

### Plant, soil, and soil slurry nutrient analysis

Composited dried and homogenized soil and plant samples were analyzed for total nitrogen (N) and carbon (C) via combustion analysis (53). Soil nitrate was measured using a flow injection analyzer (54). Soil extractable phosphorus (P) was determined according to Olsen and Page (1982) (55). Other soil exchangeable ions in soil, soil slurries and plant samples were measured using ICP-AES (56). Soil organic matter content was determined via the loss-on-ignition method (57). Soil pH was measured using a saturated paste extract.

### Rhizosphere DNA extraction and amplicon sequencing

Three plants were excavated from each plot three weeks after transplanting. Six plots were sampled per field as described above. In the lab, roots from each plant were divided into 12 subsamples. Root fragment subsamples were shaken briefly to remove adhering soil, then shaken for 90 minutes in 0.9% NaCl and 0.01% Tween80, then extracted using the MoBio PowerSoil Kit (Qiagen). At least 100 ng of rhizosphere DNA from each sample was sent for library prep and sequencing using MiSeq at Dalhousie IMR facility. The V4-V5 region of the 16SrRNA region was sequenced to characterize bacterial communities and the ITS region of the rRNA gene was sequenced to characterize fungal communities (58). Negative controls from the extraction buffer and kit materials were also submitted, but no reads were recovered. Reads were error-corrected and assembled into ASVs using DADA2 v1.8(59) and assigned taxonomy using SILVA v.128 for bacteria (60), and UNITE database (2017 release) for fungi (61). Taxa without a taxonomic assignment, or assigned to archaea, mitochondria, or chloroplasts were removed from the dataset. Those not assigned to the kingdom Fungi were removed from the fungal dataset. Sequence abundance was rarefied to 15,310 sequences per sample for bacteria and 13,000 per sample for fungi and all sampling curves approached saturation.

### Statistical Analysis

All statistics were conducted using R (R 3.2.2) (62). Assumptions of homogeneity and normal distribution of residuals were checked and data were transformed when appropriate to improve homoscedasticity or non-parametric tests were used. Wilcoxon-Mann-Whitney tests were used to determine the impact of farm management on total number of Arthropods collected in sweepnets. Student’s t-tests were performed to determine the impact of farm management on insect fecundity, survival, plant nutrition and soil properties at each paired site. The impact of management and the interaction on phytohormone levels were tested with a generalized mixed model using linear regression in lme4 with SA and JA as response variables and site as a random effect. For comparison of phytohormone levels at individual farms, linear regression was used to determine the impact of management. Insect settling data was analyzed with Binomial regression to determine the impact of soil management, chemical treatment and mutants on insect settling. Statistical differences were determined for settling assays using a binomial test assuming the null hypothesis of no difference between the treatments.

### Correlations among plant, soil, and microbial variables

Mantel tests were conducted to identify Pearson correlations among plant, soil, and microbial variables. Plant variables included shoot and root dry weight, foliar nutrient concentrations (Table S2) and log-transformed SA and JA concentrations. Soil variables included all measured soil nutrients (Table S3). Bray-Curtis dissimilarity matrices were calculated separately for bacterial and fungal ASV tables. Plots within each farm were used as replicates (*N* = 12 plots/farm pair, with *N* = 6 plots per management type within a pair). Scaled Euclidean distance matrices were calculated for plant and soil variables. Correlations between all pairs of matrices were tested for significance using permutation (mantel() function of vegan package (63). Partial Mantel tests were conducted for plant, soil, and bacterial matrices and plant, soil, and fungal matrices to determine whether plant or soil variables predicted microbial community composition if the other category of variables was held constant (mantel.partial() function of vegan package).

### Differential abundance and indicator species analysis

Bacterial and fungal OTUs whose abundance varied with SA were identified using differential abundance analysis (DESeq2 package) (64). Taxa were filtered to remove those occurring in fewer than three samples. To control for variation in SA among sites, we calculated residuals with site only as a predictor. SA residuals were log transformed and used as continuous predictors in the DESeq2 analysis. Taxa whose abundance varied significantly (*P* < .05) were identified using the Wald test. All mixed models, multivariate analyses including NMDS, RDA and mantel tests, and differential abundance analyses were conducted using R (R 3.2.2) and implemented in Rstudio (65).

## References

1. Shennan C, et al. (2017) Organic and conventional agriculture: A useful framing? Annu Rev Environ Resour 42:317–346.

2. Reganold JP, Wachter JM (2016) Organic agriculture in the twenty-first century. Nat Plants 2:15221.

3. Seufert V, Ramankutty N (2017) Many shades of gray—The context-dependent performance of organic agriculture. Sci Adv 3(3).

4. Trewavas A (2001) Urban myths of organic farming. Nature 410:409.

5. Ponisio LC, et al. (2015) Diversification practices reduce organic to conventional yield gap. Proc R Soc London B Biol Sci 282(1799).

6. Ponisio CL, Ehrlich RP (2016) Diversification, yield and a new agricultural revolution: Problems and prospects. Sustain 8: 1118.

7. Crowder DW, Northfield TD, Strand MR, Snyder WE (2010) Organic agriculture promotes evenness and natural pest control. Nature 466(7302):109–112.

8. Lichtenberg EM, et al. (2017) A global synthesis of the effects of diversified farming systems on arthropod diversity within fields and across agricultural landscapes. Glob Chang Biol 23(11):4946–4957.

9. Muneret L, et al. (2018) Evidence that organic farming promotes pest control. Nat Sustain 1(7):361–368.

10. Hole DG, et al. (2005) Does organic farming benefit biodiversity? Biol Conserv 122(1):113–130.

11. Garratt MPD, Wright DJ, Leather SR (2011) The effects of farming system and fertilisers on pests and natural enemies: A synthesis of current research. Agric Ecosyst Environ 141(3):261–270.

12. Verbruggen E, et al. (2010) Positive effects of organic farming on below-ground mutualists: large-scale comparison of mycorrhizal fungal communities in agricultural soils. New Phytol 186(4):968–979.

13. Hartmann M, Frey B, Mayer J, Mäder P, Widmer F (2015) Distinct soil microbial diversity under long-term organic and conventional farming. ISME J 9(5):1177–1194.

14. Lori M, Symnaczik S, Mäder P, De Deyn G, Gattinger A (2017) Organic farming enhances soil microbial abundance and activity—A meta-analysis and meta-regression. PLoS One 12(7)

15. Lupatini M, Korthals GW, de Hollander M, Janssens TKS, Kuramae EE (2016) Soil microbiome is more heterogeneous in organic than in conventional farming system. Front Microbiol 7:2064.

16. Gosling P, Hodge A, Goodlass G, Bending GD (2006) Arbuscular mycorrhizal fungi and organic farming. Agric Ecosyst Environ 113(1):17–35.

17. Vannette RL, Hunter MD (2009) Mycorrhizal fungi as mediators of defence against insect pests in agricultural systems. Agric For Entomol 11(4):351–358.

18. Pineda A, Zheng S-J, van Loon JJA, Pieterse CMJ, Dicke M (2010) Helping plants to deal with insects: the role of beneficial soil-borne microbes. Trends Plant Sci 15(9):507–514.

19. Berendsen RL, Pieterse CMJ, Bakker PAHM (2012) The rhizosphere microbiome and plant health. Trends Plant Sci 17(8):478–486.

20. Pozo MJ, Azcón-Aguilar C (2007) Unraveling mycorrhiza-induced resistance. Curr Opin Plant Biol 10(4):393–398.

21. Vannette RL, Hunter MD (2010) Plant defence theory re-examined: nonlinear expectations based on the costs and benefits of resource mutualisms. J Ecol 99(1):66–76.

22. Fritz M, Jakobsen I, Lyngkjær MF, Thordal-Christensen H, Pons-Kühnemann J (2006) Arbuscular mycorrhiza reduces susceptibility of tomato to *Alternaria solani*. Mycorrhiza 16(6):413–419.

23. Kempel A, Schmidt AK, Brandl R, Schädler M (2010) Support from the underground: Induced plant resistance depends on arbuscular mycorrhizal fungi. Funct Ecol 24(2):293–300.

24. Pineda A, Kaplan I, Bezemer TM (2017) Steering soil microbiomes to suppress aboveground insect pests. Trends Plant Sci 22:770–778.

25. Pangesti N, Pineda A, Pieterse CMJ, Dicke M, Van Loon JJA (2013) Two-way plant mediated interactions between root-associated microbes and insects: from ecology to mechanisms. Front Plant Sci 4:414.

26. Katayama N, Zhang ZQ, Ohgushi T (2011) Community-wide effects of below-ground rhizobia on above-ground arthropods. Ecol Entomol 36(1):43–51.

27. Chen L-F, Batuman O, Aegerter BJ, Willems J, Gilbertson RL (2017) First report of curly top disease of pepper and tomato in California caused by the spinach curly top strain of Beet curly top virus. Plant Dis 101: 1334.

28. Erb M, Meldau S, Howe GA (2012) Role of phytohormones in insect-specific plant reactions. Trends Plant Sci 17(5):250–259.

29. Bari R, Jones JD (2009) Role of plant hormones in plant defence responses. Plant Mol Biol 69(4):473–488.

30. Prudic KL, Oliver JC, Bowers MD (2005) Soil nutrient effects on oviposition preference, larval performance, and chemical defense of a specialist insect herbivore. Oecologia 143: 578–587.

31. Lu Z, Yu X, Heong K, Hu C (2007) Effect of nitrogen fertilizer on herbivores and its stimulation to major insect pests in rice. Rice Sci 14: 56–66.

32. Schmidt J, Vannette RL, Igwe A, Blundell B, Casteel CL, Gaudin (2019) Effects of agricultural management on rhizosphere microbial structure and function in processing tomato. Appl Environ Microbiol 85 (16): e01064–1.

33. Pieterse CMJ, et al. (2014) Induced systemic resistance by beneficial microbes. Annu Rev Phytopathol 52: 347–375.

34. Wolf KM, et al. (2018) The century experiment: the first twenty years of UC Davis’ mediterranean agroecological experiment. Ecology 99: 503.

35. Brading PA, Hammond-Kosack KE, Parr A, Jones JDG (2000) Salicylic acid is not required for Cf-2- and Cf-9-dependent resistance of tomato to *Cladosporium fulvum*. Plant J 23: 305–318.

36. Walling LL (2008) Avoiding effective defenses: Strategies employed by phloem-feeding insects. Plant Physiol 146:859–866.

37. Kaloshian I, Walling LL (2005) Hemipterans as plant pathogens. Annu Rev Phytopathol 43:491–521.

38. Rodríguez-Álvarez CI, López-Climent MF, Gómez-Cadenas A, Kaloshian I, Nombela G (2015) Salicylic acid is required for Mi-1-mediated resistance of tomato to whitefly *Bemisia tabaci*, but not for basal defense to this insect pest. Bull Entomol Res 105:574–582.

39. Ellis C, Karafyllidis L, Turner JG (2002) Constitutive activation of jasmonate signaling in an Arabidopsis mutant correlates with enhanced resistance to Erysiphe cichoracearum, Pseudomonas syringae, and Myzus persicae. Mol Plant-Microbe Interact 15(10):1025–1030.

40. Kloth KJ, et al. (2016) AtWRKY22 promotes susceptibility to aphids and modulates salicylic acid and jasmonic acid signalling. J Exp Bot 67(11):3383–3396.

41. Cui J, et al. (2005) *Pseudomonas syringae* manipulates systemic plant defenses against pathogens and herbivores. Proc Natl Acad Sci 102: 1791–1796.

42. De Souza R, Ambrosini A, Passaglia LMP (2015) Plant growth-promoting bacteria as inoculants in agricultural soils. Genet Mol Biol 4: 401–419.

43. Nakano M, Mukaihara T (2018) *Ralstonia solanacearum* type III effector RipAL targets chloroplasts and induces jasmonic acid production to suppress salicylic acid-mediated defense responses in plants. Plant Cell Physiol 59: 2576–2589.

44. Baichoo Z, Jaufeerally-Fakim Y (2017) *Ralstonia solanacearum* upregulates marker genes of the salicylic acid and ethylene signaling pathways but not those of the jasmonic acid pathway in leaflets of Solanum lines during early stage of infection. Eur J Plant Pathol 147: 615–625.

45. Barber NA, Kiers ET, Theis N, Hazzard R V., Adler LS (2013) Linking agricultural practices, mycorrhizal fungi, and traits mediating plant-insect interactions. Ecol Appl 23(7):1519–1530.

46. Berg M, Koskella B (2018) Nutrient- and dose-dependent microbiome-mediated protection against a plant pathogen. Curr Biol 28:2487–2492.

47. Blubaugh CK, Carpenter-Boggs L, Reganold JP, Schaeffer RN, Snyder WE (2018) Bacteria and competing herbivores weaken top–down and bottom–up aphid suppression. Front Plant Sci 9

48. Heinen R, et al. (2018) Species-specific plant–soil feedbacks alter herbivore-induced gene expression and defense chemistry in *Plantago lanceolata*. Oecologia 188(3):801–811.

49. Bastías DA, et al. (2018) Jasmonic acid regulation of the anti-herbivory mechanism conferred by fungal endophytes in grasses. J Ecol 106(6):2365–2379.

50. Gilbert L, Johnson D (2015) Plant-mediated “apparent effects” between mycorrhiza and insect herbivores. Curr Opin Plant Biol 26(Figure 2):100–105.

## References

51. Lightner, J, Pearce G, Ryan CA, Browse J (1993) Isoaltion of signalling mutants of tomato (*Lycopersicon esculentum*). Mol Genet Genomics 241: 595–601.

52. Casteel CL, et al. (2015) Disruption of ethylene responses by Turnip mosaic virus mediates suppression of plant defense against the green peach aphid vector. Plant Physiol 169: 209–218.

53. AOAC Official Method 972.43, Microchemical determination of carbon, hydrogen, and nitrogen, automated method (2006). Official Methods of Analysis of AOAC International (AOAC International, Gaithersburg, MD), pp 5–6. 18th Ed.

54. Knepel K (2003) Determination of nitrate in 2M KCl soil extracts by flow injection analysis. QuikChem Method 12(107).

55. Olsen SR, Page AL (1982) Phosphorus. Methods of soil analysis. Part 2. Chemical and Microbiological Properties. Agron. Monogr. 9. doi:10.2134/agronmonogr9.2.2ed.

56. Jones JB (2001) Laboratory guide for conducting soil tests and plant analysis.

57. Nelson DW, Sommers LE (1996) Total carbon, organic carbon, and organic matter.

58. Walters W, et al. (2016) Improved bacterial 16S rRNA gene (V4 and V4-5) and fungal internal transcribed spacer marker gene primers for microbial community surveys. mSystems 1(1).

59. Callahan BJ, et al. (2016) DADA2: High-resolution sample inference from Illumina amplicon data. Nat Methods 13: 581–583.

60. Glöckner FO, et al. (2017) 25 years of serving the community with ribosomal RNA gene reference databases and tools. J Biotechnol 261: 169–176.

61. Kõljalg U, et al. (2013) Towards a unified paradigm for sequence-based identification of fungi. Mol Ecol 22: 5271–5277.

62. R Core Team (2014) R: A language and environment for statistical computing.

63. Oksanen, J. Blanchet, F.G. Friendly, M. Kindt, P.L. McGlinn, D. Minchin, P.R., O’Hara, R.B. Simpson, G.K. Solymos, P. Stevens, H.H. Szoecs, E. Wagner H (2018) vegan: Community Ecology Package.

64. Love MI, Huber W, Anders S (2014) Moderated estimation of fold change and dispersion for RNA-seq data with DESeq2. Genome Biol 15: 550–571.

65. RStudio Team (2015) RStudio: Integrated development for R.

